# Efficient inference of population size histories and locus-specific mutation rates from large-sample genomic variation data

**DOI:** 10.1101/006742

**Authors:** Anand Bhaskar, Y.X. Rachel Wang, Yun S. Song

## Abstract

With the recent increase in study sample sizes in human genetics, there has been growing interest in inferring historical population demography from genomic variation data. Here, we present an efficient inference method that can scale up to very large samples, with tens or hundreds of thousands of individuals. Specifically, by utilizing analytic results on the expected frequency spectrum under the coalescent and by leveraging the technique of automatic differentiation, which allows us to compute gradients exactly, we develop a very efficient algorithm to infer piecewise-exponential models of the historical effective population size from the distribution of sample allele frequencies. Our method is orders of magnitude faster than previous demographic inference methods based on the frequency spectrum. In addition to inferring demography, our method can also accurately estimate locus-specific mutation rates. We perform extensive validation of our method on simulated data and show that it can accurately infer multiple recent epochs of rapid exponential growth, a signal which is difficult to pick up with small sample sizes. Lastly, we apply our method to analyze data from recent sequencing studies, including a large-sample exome-sequencing dataset of tens of thousands of individuals assayed at a few hundred genic regions.

**Software:** We have implemented the algorithms described in this paper in an open-source software package called fastNeutrino, which stands for **fast Ne** (effective population size) and m**ut**ation **r**ate **i**nference using a**n**alytic **o**ptimization. It will be publicly available at https://sourceforge.net/projects/fastneutrino.

## Introduction

A commonly used null model in population genetics assumes that individuals are randomly sampled from a well-mixed population of constant size that evolves neutrally according to some model of random mating (Ewens 2004). However, several recent large-sample sequencing studies in humans (Coventry et al. 2010; Nelson et al. 2012; Tennessen et al. 2012; Fu et al. 2012) have found an excess of single nucleotide variants (SNVs) that have very low minor allele frequency (MAF) in the sample compared to that predicted by coalescent models with a constant effective population size. For example, in a sample of approximately 12,500 individuals of European descent analyzed by Nelson et al. (2012), over 74% of the SNVs have only one or two copies of the minor allele, and that over 95% of the SNVs have MAF less than 0.5%. On the other hand, assuming a constant population size over time, Kingman’s coalescent predicts that the number of neutral SNVs is inversely proportional to the sample frequency of the variant (Fu 1995). Keinan and Clark (2012) have suggested that such an excess of sites segregating with low MAF can be explained by recent exponential population growth. In particular, a rapid population expansion produces genealogical trees which have long branch lengths at the tips of the trees, leading to a large fraction of mutations being private to a single individual in the sample. Motivated by these findings and rapidly increasing sample sizes in population genomics, we here tackle the problem of developing an efficient algorithm for inferring historical effective population sizes and locus-specific mutation rates using a very large sample, with tens or hundreds of thousands of individuals.

At the coarsest level, previous approaches to inferring demography from genomic variation data can be divided according to the representation of the data that they operate on. Full sequence-based approaches for inferring the historical population size such as the works of Li and Durbin (2011) and Sheehan et al. (2013) use between two to a dozen genomes to infer piecewise constant models of historical population sizes. Since these approaches operate genome wide, they can take into account linkage information between neighboring SNVs. On the other hand, they are computationally very expensive and cannot be easily applied to infer recent demographic events from large numbers of whole genomes. A slightly more tractable approach to inferring potentially complex demographies involves comparing the length distribution of identical-by-descent and identical-by-state tracts between pairs of sequences (Palamara et al. 2012; Harris and Nielsen 2013).

The third class of methods, and the one that our approach also belongs to, summarizes the variation in the genome sequences by the sample frequency spectrum (SFS). The SFS of a sample of size *n* counts the number of SNVs as a function of their mutant allele frequency in the sample. Since the SFS is a very efficient dimensional reduction of large-scale population genomic data that summarizes the variation in *n* sequences by *n −* 1 numbers, it is naturally attractive for computational and statistical purposes. Furthermore, the expected SFS of a random sample drawn from the population strongly depends on the underlying demography, and there have been several previous approaches that exploit this relationship for demographic inference. Nielsen (2000) developed a method based on coalescent tree simulations to infer exponential population growth from single nucleotide polymorphisms that are far enough apart to be in linkage equilibrium. Coventry et al. (2010) developed a similar coalescent simulation-based method that additionally infers per-locus mutation rates, and applied this method to exome-sequencing data from approximately 10,000 individuals at 2 genes. Nelson et al. (2012) have also applied this method to a larger dataset of 11,000 individuals of European ancestry (CEU) sequenced at 185 genes to infer a recent epoch of exponential population growth. The common feature of all these methods is that they use Monte Carlo simulations to empirically estimate the expected SFS under a given demographic model, and then compute a pseudo-likelihood function for the demographic model by comparing the expected and observed SFS. The optimization over the demographic models is then performed via grid search procedures. More recently, Excoffier et al. (2013) have developed a software package that employs coalescent tree simulations to estimate the expected joint SFS of multiple subpopulations for inferring potentially very complex demographic scenarios from multi-population genomic data. The problem of demographic inference has also been approached from the perspective of diffusion processes. Given a demographic model, one can derive a partial differential equation (PDE) for the density of segregating sites at a given derived allele frequency as a function of time. Gutenkunst et al. (2009) used numerical methods to approximate the solution to this PDE, while Lukić et al. (2011) approximated this solution using an orthogonal polynomial expansion.

### The main novelties of our method

Similar to the inference methods of Nielsen, Coventry et al. and Excoffier et al., we also work in the coalescent framework rather than in the diffusion setting. However, our method differs from existing methods in several ways. First, our method is based on an efficient algorithmic adaption of the analytic theory of the expected SFS for deterministically varying population size models that was developed by Polanski et al. (2003) and Polanski and Kimmel (2003). This is in contrast to expensive Monte-Carlo coalescent simulations employed by the aforementioned coalescent-based methods. As a result, our approach is much more efficient and allows us to more thoroughly search the space of demographic models of interest. In a related work, Marth et al. (2004) developed analytic expressions for the expected SFS of *piecewise-constant* population size models, and used them for inferring piecewise-constant population size histories for European, Asian and African-American populations. However, their analytic expressions are not numerically stable for sample sizes larger than about 50 individuals, and, moreover, do not extend to more general population size models. On the other hand, our approach utilizes numerically stable expressions for the expected SFS developed by Polanski and Kimmel (2003) which extend to *arbitrary* variable population size models.

Second, our method uses the Poisson Random Field (PRF) approximation proposed by Sawyer and Hartl (1992). Under this approximation, the segregating sites within a locus are assumed to be far enough apart to be completely unlinked. This is also the same approximation made by numerical and spectral methods for inference under the Wright-Fisher diffusion process (Gutenkunst et al. 2009; Lukić et al. 2011), and by the coalescent method of Excoffier et al. At the other extreme, the method of Coventry et al. assumes that all the segregating sites in a locus are completely linked and hence share the same underlying genealogy. Both these model simplifications — the assumption of perfectly linked or completely independently evolving sites within a locus — are biologically unrealistic. However, as we demonstrate in our results on simulated data, our method can recover demographic parameters accurately even when the data are generated under realistic recombination rates that are inferred from human genetics studies. Working in the PRF model also confers our method a significant computational benefit. Under this assumption, we can derive efficiently computable expressions for the maximum likelihood estimate of the mutation rates at each locus. This contrasts with coalescent simulation-based methods where either the mutation rate is assumed to be known (Excoffier et al. 2013), or a grid search has to be performed over the mutation rates (Coventry et al. 2010; Nelson et al. 2012). This makes our method orders of magnitude more efficient.

Third, our method has advantages over the diffusion-based methods of Gutenkunst et al. (2009) and Lukić et al. (2011). By working in the coalescent framework, the running time of our method is independent of the population size. Furthermore, by leveraging analytic and numerically stable results for the expected SFS under the coalescent, our computations are exact. On the other hand, the method of Gutenkunst et al. (2009) must carefully discretize the allele frequency space to minimize the accumulation of numerical errors, while the method of Lukić et al. (2011) has to choose a suitable order for the spectral expansion of the transition density function of the diffusion process.

Finally, since our method is based on a likelihood function that is computed exactly, we can take advantage of the technique of automatic differentiation (Griewank and Corliss 1991) to compute exact gradients of the likelihood function. This is one of the key novel features of our approach, and obviates the need for doing a grid search over the parameters. Instead, we take advantage of efficient gradient-based algorithms for optimization over the space of demographic parameters.

### Statistical identifiability

Before one can perform any meaningful demographic inference from SFS data, it is worth examining whether the statistical problem is well defined. Myers et al. (2008) showed that, in general, the historical effective population size function is not uniquely determined by the SFS, no matter how large the sample size considered. In particular, they showed that there are infinitely many population size functions that can generate the same expected SFS for all sample sizes. While this non-identifiability result might seem like a barrier to statistical inference, it was recently shown (Bhaskar and Song 2013) that if the amount of fluctuation in the effective population size is bounded, then the expected SFS of even a moderate-size sample can uniquely identify the underlying population size function. In this work, we perform inference under the assumption that the true population size function is piecewise defined, where each piece has either a constant or exponentially growing/declining size. Since our method relies on analytic theory for the expected SFS that extends to any variable population size model, it can be easily extended to perform inference under any parametric demographic model that is identifiable from the expected SFS of a finite sample; general bounds on the sample size sufficient for identifiability were obtained by Bhaskar and Song (2013).

### Software

We have implemented the algorithms described in this paper in an open-source software package called fastNeutrino, which stands for **fast Ne** (effective population size) and m**ut**ation **r**ate **i**nference using a**n**alytic **o**ptimization. It will be publicly available at https://sourceforge.net/projects/fastneutrino.

## Method

In this section, we provide an overview of our method. More involved computational details are provided in the Supplemental material. In our work we utilize automatic differentiation (Griewank and Corliss 1991), a technique that allows one to compute exact gradients numerically without requiring explicit symbolic expressions for the gradient. This technique has not been widely adopted in population genetics, so we include a brief exposition of it here.

### Model and Notation

We assume that our data are drawn according to Kingman’s coalescent from a single panmictic population having population size *N* (*t*) haploids at time *t*, where *t* is increasing in the past. Without loss of generality, we assume that the sample is drawn from the population at the present time *t* = 0. We shall also assume the infinite-sites model of mutation, where mutations occur at a low enough rate that any segregating site in the sample has experienced at most one mutation event.

The data we wish to analyze, denoted by 𝒟, consist of the sample frequency spectrum (SFS) for *n* haploid (or *n/*2 diploid) individuals at each of *L* loci located sufficiently far apart along the genome. The SFS at locus *l* is a vector 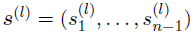, where 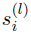 is the number of segregating sites which have *i* copies of the mutant allele among the *n* alleles at that site. We are also given the length *m*^(*l*)^ of each locus *l*. For notational convenience, let 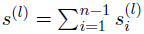 be the total number of segregating sites in the sample at locus *l*. Given the data 𝒟, our goal is to infer the haploid effective population size function *N* (*t*) and the per-base locus-specific mutation rates *µ*^(*l*)^. We use Φ to denote a vector of parameters that parameterize the family of piecewise-exponential demographic models. Note that such a family also contains piecewise-constant population size functions. While we describe our method assuming knowledge of the identities of the ancestral and mutant alleles, we can just as easily work with the folded SFS which counts the segregating sites as a function of the sample minor allele frequency, if the identity of the ancestral allele is not known.

### Likelihood

Let us first restrict attention to a single locus *l*. For a locus with length *m* bases and per-base per-generation mutation rate *µ*, let *θ/*2 denote the population-scaled mutation rate for the whole locus. Specifically,

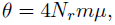

where *N_r_* denotes a reference population size which is used as a scaling parameter.

We wish to compute the probability of the observed frequency spectrum ***s*** = (*s*_1_*,…, s_n−_*_1_) at locus *l* under the infinite-sites model. (We omit the superscript *l* for ease of notation.) If all the sites in the locus are completely linked and the *n* individuals in the sample are related according to the coalescent tree *T,* then the probability of observing the frequency spectrum ***s*** is given by

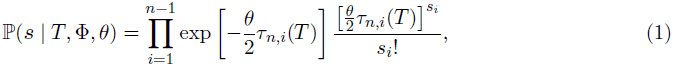

where *τ_n,i_*(*T)* is the sum of the lengths of branches in the coalescent tree *T* which subtend *i* descendant leaves. The explanation for (1) is as follows: in the infinite-sites mutation model, mutations occur on the coalescent tree according to a Poisson process with rate *θ*/2, where every mutation generates a new segregating site. A mutation creates a segregating site with *i* mutant alleles if and only if it occurs on a branch that subtends *i* descendants in the sample. To avoid unwieldy notation, we drop the dependence on the tree *T* for the branch lengths *τ_n,i_*(*T)*. To compute the probability of the observed frequency spectrum ***s***, we need to integrate (1) over the distribution *f* (*T |* Φ) of *n*-leaved coalescent trees *T* under the demography Φ. Let *𝒯_n_* denote the space of coalescent trees with *n* leaves. Then, abusing notation, the probability ℙ(***s*** | Φ*, θ*) can be written as

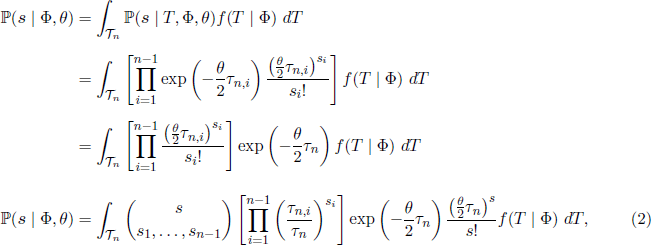

where 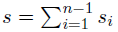. In (2), 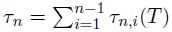, the total branch length of the tree *T* on *n* haploid individuals. It is not known how to efficiently and exactly compute (2), even when Φ represents the constant population size demographic model. Most works approximate the integral in (2) by sampling coalescent trees under the demographic model Φ. In order to find the MLE for *θ*, they must repeat this Monte-Carlo integration for each value of *θ* in some grid.

### Poisson Random Field approximation

In our method, we use the Poisson Random Field (PRF) assumption of Sawyer and Hartl (1992) which assumes that all the sites in a given locus are completely unlinked, and hence the underlying coalescent tree at each site is independent. Under this assumption, the probability of the observed frequency spectrum ***s*** is given by

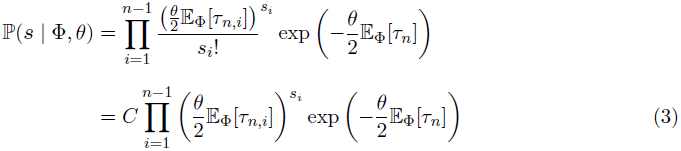

where the expectations 𝔼_Φ_[·] in (3) are taken over the distribution on coalescent trees with *n* leaves drawn from the demographic model parameterized by Φ, and 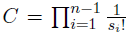 is a data-dependent constant that can be ignored for maximum likelihood estimation.

Hence, under the PRF approximation, the problem of computing the likelihood in (3) reduces to that of computing the expectations 𝔼_Φ_[*τ_n,i_*] and 𝔼_Φ_[*τ_n_*] for the demographic model given by Φ. Using analytic results for the SFS for variable population sizes developed by Polanski et al. (2003) and Polanski and Kimmel (2003), we can develop an efficient algorithm to *numerically stably* and *exactly* compute 𝔼_Φ_[*τ_n,i_*] and 𝔼_Φ_[*τ_n_*] for a wide class of population size functions *N* (*t*). In this work, we consider inference in the family of piecewise-exponential functions with either a prescribed or variable number of pieces. The details of computing 𝔼_Φ_[*τ_n,i_*] and 𝔼_Φ_[*τ_n_*] for such a class of population size functions is given in the Supplemental material. Inference under other families of parametric demographic models can just as easily be performed if one can efficiently compute the integral expressions given in the Supplemental material.

Taking logarithms on both sides in (3), we get the following log-likelihood for the demographic model Φ and mutation rate *θ* at this locus:

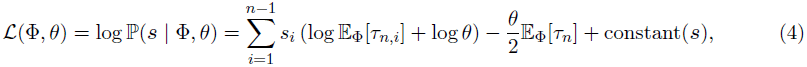

where constant(***s***) depends on ***s*** but not on the parameters Φ*, θ*.

Assuming the loci are all completely unlinked, the log-likelihood for one locus given in (4) can be summed across all loci *l* = 1*,…, L* to get a log-likelihood for the entire dataset 𝒟:

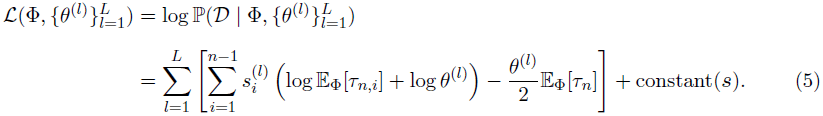

It is easy to see that *ℒ* is a concave function of the mutation rates *θ*^(*l*)^, since the Hessian *H* of *ℒ* with respect to ***θ*** = (*θ*^(1)^*,…, θ*^(*L*)^) is given by

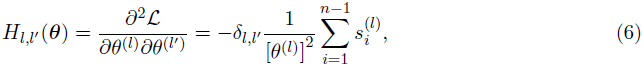

showing that *H*(***θ***) is negative definite for all ***θ*** ≻ **0**. Hence, the mutation rates of the loci that maximize ℒ are the solutions of

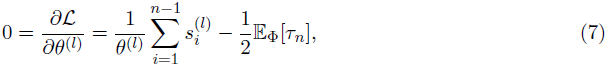

yielding the following maximum likelihood estimate for the mutation rate *θ*^(*l*)^ at locus *l* given the demographic model:

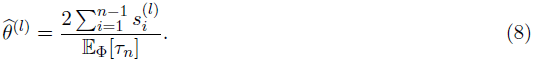

Note that for a constant population size, (8) is the same as Watterson’s estimator 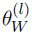 for the mutation rate (Watterson 1975), namely

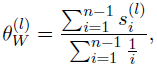

since for a constant population size, 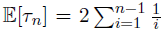. Substituting the MLE for *θ*^(*l*)^ in (8) into (5), we obtain the log-likelihood with the optimal mutation rates:

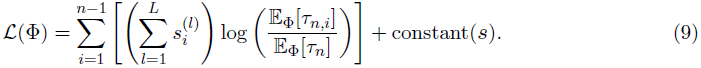

If we define the discrete probability distributions 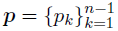 and 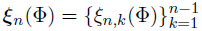 by

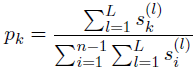

and

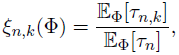

then we see that the demographic model Φ̭ that is the MLE of the likelihood function *ℒ*(Φ) in (9) is given by

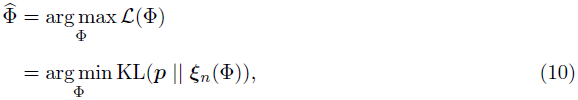

where KL(*P || Q*) denotes the Kullback-Liebler divergence of distribution *Q* from *P*.

Hence, for a given demographic model Φ, we can compute the log-likelihood using (9), and infer the optimal mutation rate at each locus independently according to (8). We compute the gradient of *ℒ*(Φ) with respect to Φ using automatic differentiation (Griewank and Corliss 1991), detailed below. Once we have access to the gradient of *ℒ*(Φ), we can more efficiently search over the space of demographic models using standard gradient-based optimization algorithms.

### Some computational details

For a sample of size *n* and a piecewise-exponential demographic model with *M* epochs, the terms 𝔼_Φ_[*τ_n,i_*] in (9) can be computed in *O*(*nM)* time for each value of the index *i*, 1 *≤ i ≤ n −* 1. Hence, the time complexity for evaluating the log-likelihood function in (9) is *O*(*n*^2^ *M)*. However, since we would like to use our method for large sample sizes on the order of tens to hundreds of thousands of individuals, in practice, and for the results reported in **Results**, we evaluate (9) by using only the leading entries of the SFS which account for some significant fraction of the segregating sites in the observed sample. In particular, for the results on simulated and real datasets reported in **Results**, we use the first *k* entries of the SFS which account for 90% of the segregating sites in the observed data and collapse the remaining *n* − *k* − 1 SFS entries into one class when computing the KL divergence in (10).

### Confidence intervals

If we ignore the fact that we are using the PRF assumption to treat the sites within each locus as freely recombining, (9) is the log-likelihood function of *L* independent samples from a multinomial distribution with *n −* 1 categories and probabilities 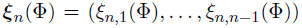, where 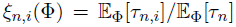, and 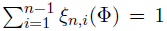. Since the probability models specified by the likelihood function *ℒ*(Φ) are identifiable for a sufficiently large sample size *n* that depends on the number of parameters being inferred (Bhaskar and Song 2013), and since ***ξ****_n_*(Φ) is differentiable with respect to Φ, it follows from the asymptotics of maximum likelihood estimators that

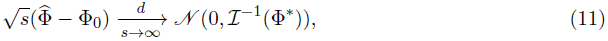

where Φ* is the true underlying demographic model, and ℐ(Φ*) is the expected Fisher information matrix of a single observation. The elements of ℐ(Φ*) are given by

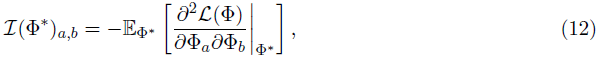

where Φ*_i_* denotes the *i*th element of the demographic parameter vector Φ. For the log-likelihood function *ℒ*(Φ) in (9), using 𝔼_Φ*\**_ (*p_k_*) = *ξ_n,k_*(Φ*^*^*), it is straightforward to show that (12) simplifies to

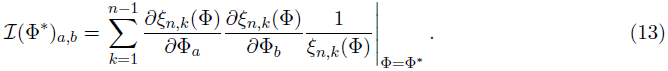

We calculate the partial derivatives in (13) via automatic differentiation during the computation of ***p***(Φ) (Griewank and Corliss 1991). This allows us to construct asymptotic empirical confidence intervals for Φ*. An asymptotic 100(1 *− α*)% confidence interval for the parameters of Φ* is given by

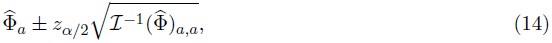

where *z_α/_*_2_ is the 100(1 *− α/*2)th percentile of a standard normal distribution.

If the loci being analyzed are long with low levels of intralocus recombination, then (9) is a composite log-likelihood function rather than a true likelihood function. In such cases, we compute empirical confidence intervals for the demographic parameters using a nonparametric block bootstrap procedure (Efron and Tibshirani 1986). In particular, we subsample *L* loci with replacement from the original dataset of *L* loci, where each locus is sampled with probability proportional to the number of segregating sites in it. We examine the performance of the asymptotic confidence interval procedure and the block bootstrap on simulated data in **Results**.

### Automatic differentiation

Since our inference algorithm and asymptotic confidence interval estimation procedure rely heavily on computing gradients via automatic differentiation, we briefly describe this technique here to keep the paper self-contained. Automatic differentiation (AD) is a powerful technique for computing the gradient (and higher-order derivatives) of a mathematical function that is implemented in a computer program. The basic idea in AD is to track the values and gradients of the intermediate variables in a computer program evaluated at the desired value of the independent variables, where the chain rule of calculus is repeatedly employed to compute the gradients. For example, suppose we had the mathematical function *f* (*x*) = sin(*x*^2^ + *x*) that is implemented via the following computer function f(x):

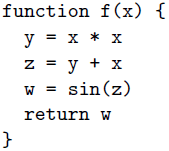

To calculate the numerical value of *df /dx*, we augment the intermediate variables in this program to create a new variable per existing variable that will track the derivative with respect to *x*, evaluated at the value passed for the argument *x*. In particular, in this program, we define *x′* = *dx/dx* (which will be trivially initialized to 1), *y′* = *dy/dx*, *z′* = *dz/dx*, and *w′* = *dw/dx*. Suppose we wish to evaluate *f* and its gradient (with respect to *x*) at *x* = 1*/*2. The desired *df /dx* evaluated at *x* = 1*/*2 is equal to *w′* evaluated at *x* = 1*/*2. The key idea in AD is to evaluate *y′, z′*, and *w′* at the same time that *y, z*, and *w* are being evaluated for the input variable value *x* = 1*/*2. These augmented variables *y′, z′*, and *w′* can be computed using the chain rule of calculus. The evaluation of the original and augmented variables of the function at *x* = 1*/*2 proceeds as follows:

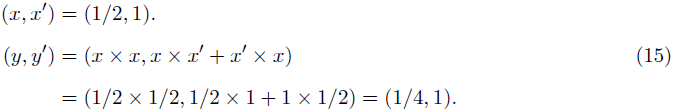

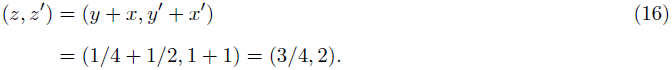

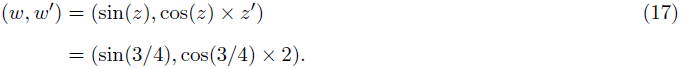

Note that the chain rule was invoked in (15), (16), and (17) to calculate *y′, z′*, and *w′* at *x* = 1*/*2. Using features of modern programming languages such as function and operator overloading, the chain rule calculations for common mathematical functions can be abstracted away in a library, thus requiring minimal changes to the user’s program. In our demographic inference software package, we used the AD library ADOL-C (Walther and Griewank 2012).

AD offers advantages over both symbolic and numerical methods for gradient evaluation. Since one does not have to derive and implement potentially complicated expressions for the symbolic gradient of the original mathematical function, AD helps reduce the chances of implementation errors. At the same time, AD computes *exact* gradients as opposed to approximate numerical methods like finite-difference schemes. The description provided above is called forward-mode AD; there are also other evaluation orders for the terms in the chain rule computation in AD. We refer the interested reader to Griewank and Corliss (1991) for a more detailed survey of this topic.

## Results

In this section, we perform extensive validation of our method on simulated data using several sets of demographic models similar to those inferred by recent large-sample studies. We also apply our method to the neutral region dataset of Gazave et al. (2014) and the exome-sequencing dataset of Nelson et al. (2012) to detect signatures of recent exponential population growth.

### Simulated demographic models

To validate our inference algorithm, we simulated data under the coalescent with recombination with the following two demographic scenarios:

- Scenario 1 — This demographic scenario models two ancestral population bottlenecks followed by an epoch of exponential growth. We simulated datasets with several values for the growth duration *t*_1_ and per-generation growth rate *r*_1_ such that the population expansion factor (1 + *r*_1_)*^t^*^1^ is fixed at 512, which is close to the estimated population expansion factor inferred by Nelson et al. (2012) in the CEU subpopulation. The ancestral population bottlenecks reflect the out-of-Africa bottleneck and the European-Asian population split, and these parameters were set to those estimated by Keinan et al. (2007). The population size functions for this scenario are shown in Figure 1(a).
- Scenario 2 — In this scenario, there are two epochs of exponential growth, the older of which (called epoch 2) lasts for *t*_2_ = 300 generations with a growth rate of *r*_2_ = 1% per generation, and a recent epoch (called epoch 1) of more rapid growth lasting *t*_1_ = 100 generations with a growth rate of *r*_1_ = 4% per generation. This model also incorporates the two ancestral population bottlenecks inferred by Keinan et al. (2007), and is shown in Figure 1(b).

**Figure 1.**
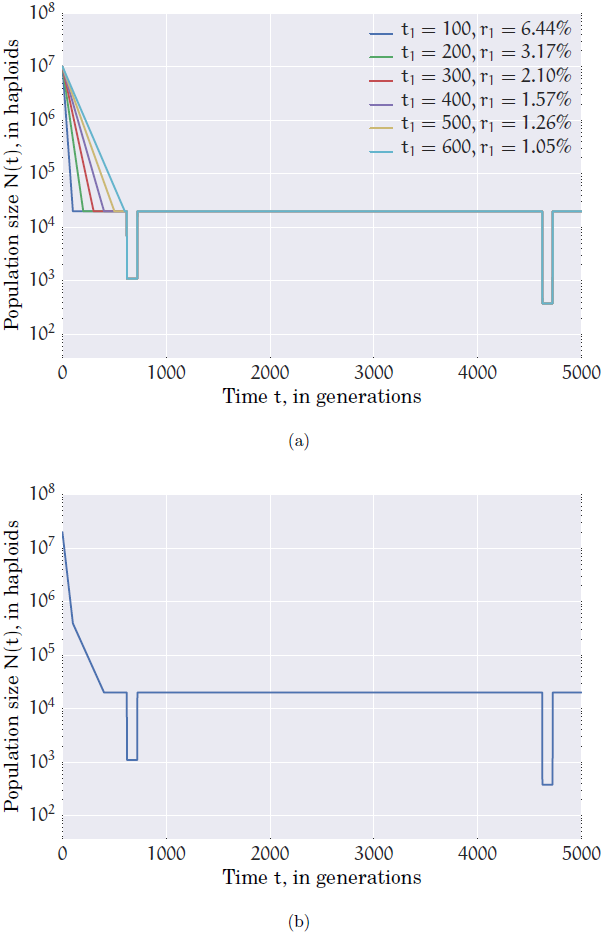
Population size *N* (*t*) as a function of time (measured in generations) for (a) several choices of *t*_1_ and *r*_1_ in Scenario 1, and (b) Scenario 2. The present time corresponds to *t* = 0.

For each demographic scenario, we simulated several datasets using the coalescent simulator ms (Hudson 2002) with 10,000 diploid individuals and 100 unlinked loci each of length 10 kb, while using a realistic recombination rate of 10^−8^ per base per haploid per generation within each locus. We used a mutation rate of 2.5 *×* 10^−8^ per base per haploid per generation at each locus.

### Maximum likelihood estimates

We applied our method to estimate the exponential growth rate and onset times for Scenario 1 and Scenario 2 while assuming that the details of the ancestral population bottlenecks is known. We did not try to estimate the ancestral population bottlenecks because our focus was on inferring recent population expansion events which are not detectable with small sample sizes. To infer ancient demographic events such as bottlenecks, we think that genome-wide methods such as those of Li and Durbin (2011) and Sheehan et al. (2013) will be more powerful. Figure 2 shows violin plots of the inferred values of the duration and rate of exponential growth for each of the simulation parameter settings in Scenario 1, with the joint distribution over the inferred parameters shown in Figure 3. In each simulation parameter combination in Scenario 1, the population expands by a factor of 512 in the epoch of recent exponential growth. The solid red curves in Figure 3 represent the exponential growth parameter combinations that have this same population expansion factor, while the dashed red curves are the parameter combinations having 25% higher and lower population expansion factors. As can be seen from the tight clustering of points along the red curves in Figure 3, the jointly inferred exponential growth parameter combinations quite accurately reflect the exponential population expansion factor — i.e. when the inferred growth onset time is large, the inferred growth rate is correspondingly lower so that the population expands by the same factor in the inferred demographic model as in the true demographic model. Similarly, Figure 4 shows the marginal distribution of the inferred values of the growth onset times and rates for each of the two epochs in Scenario 2, with the joint distribution of the inferred parameters for each of the two epochs shown in Figure 5. Since most points in Figure 5 fall within the dashed red curves, this indicates that the population expansion factors from the inferred estimates match the true population expansion factor in each epoch quite well.

**Figure 2.**
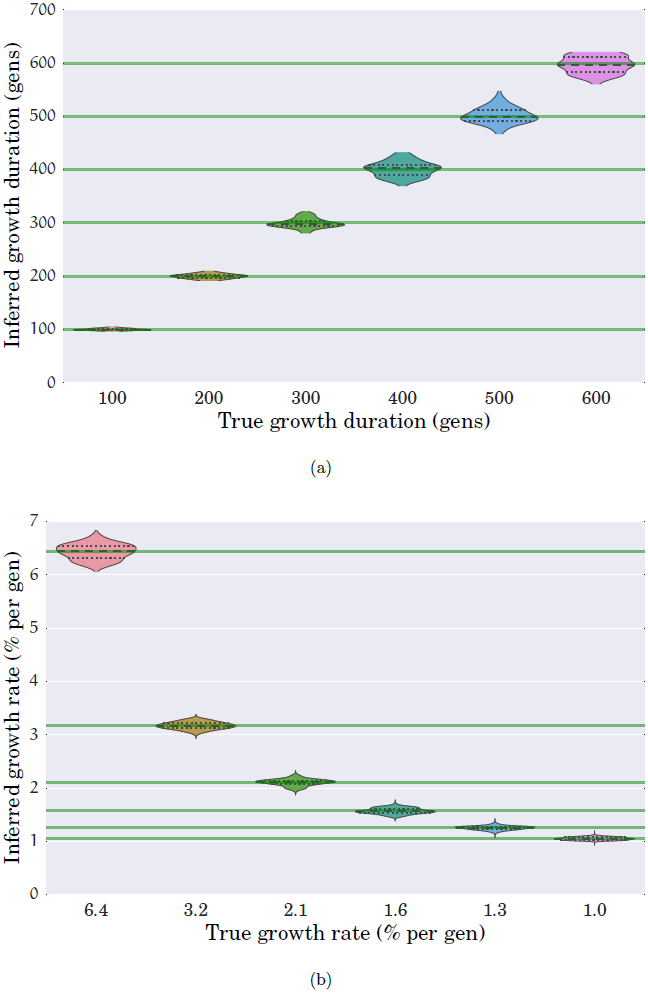
Violin plots of the (a) duration and (b) rate of exponential growth in the population size for each of the six simulation parameter settings of Scenario 1. Each violin plot is generated using 100 simulated datasets with 100 unlinked loci of 10 kb each over 10,000 diploid individuals. The green lines indicate the true values for the simulation parameters. The median inferred parameter values match the true parameter values very well.

**Figure 3.**
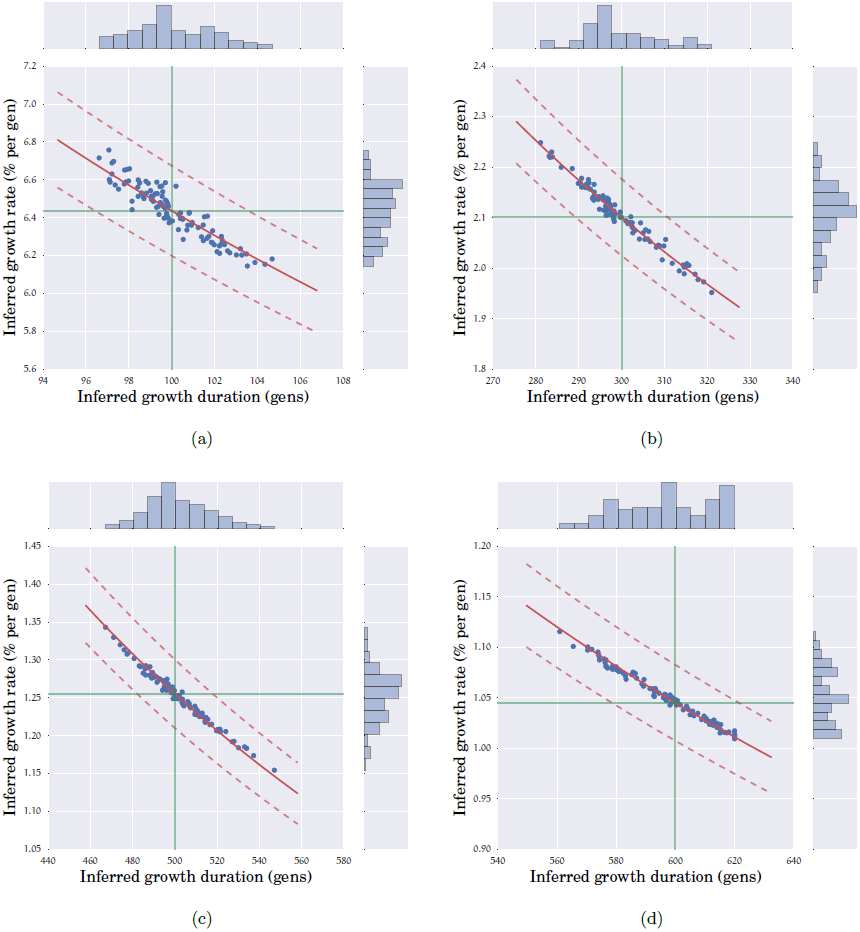
Joint and marginal distributions of the inferred duration and rate of exponential growth for several parameter settings in Scenario 1. Each point in each plot represents a simulated dataset with 100 unlinked loci of length 10 kb each over 10,000 diploid individuals. The green vertical and horizontal lines denote the true simulation parameter values for the duration and rate of exponential population growth. In Scenario 1, the present population size is 512 times larger than the ancestral population size. The solid red curves (dashed red curves) in each plot are the combinations of parameter values which have this same ratio (resp., higher and lower by 25%) of present to ancestral population size. (a) *t*_1_ = 100 gens, *r*_1_ = 6.44% per gen, (b) *t*_1_ = 300 gens, *r*_1_ = 2.10% per gen, (c) *t*_1_ = 500 gens, *r*_1_ = 1.26% per gen, (d) *t*_1_ = 600 gens, *r*_1_ = 1.05% per gen.

**Figure 4.**
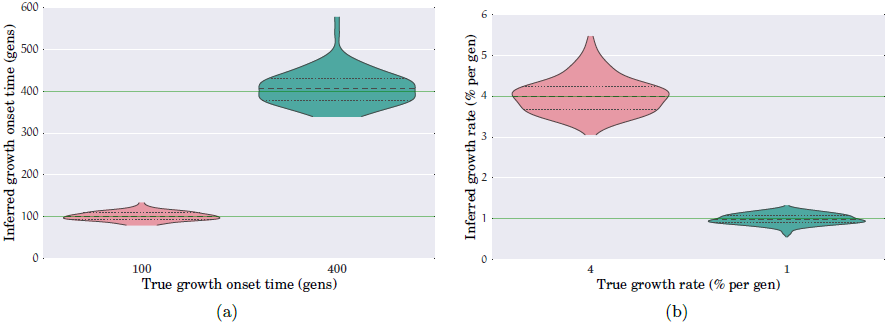
Violin plots of the (a) onset times *t*_1_ and *t*_2_, and (b) exponential growth rates *r*_1_ and *r*_2_ for the two epochs of exponential growth in Scenario 2 (see Figure 1(b)). Each violin plot represents 100 simulated datasets with 100 unlinked loci of 10 kb each over 10,000 diploid individuals. The green lines indicate the true values for the simulation parameters.

**Figure 5.**
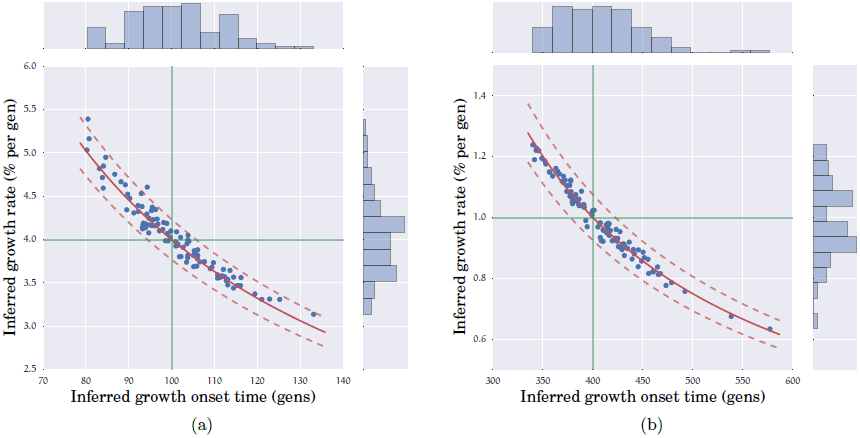
Joint distribution of the inferred growth onset time and growth rate in Scenario 2 for (a) epoch 1 with *t*_1_ = 100 gens and *r*_1_ = 4% per gen, and for (b) epoch 2 with *t*_2_ = 400 gens and *r*_2_ = 1% per gen. These plots were generated using 100 simulated datasets with 100 unlinked loci of 10 kb each over 10,000 diploid individuals. The green lines indicate the true values for the simulation parameters. In Scenario 2, the population expands about 50-fold and 20-fold in epochs 1 and 2, respectively. The solid red curves are the combinations of parameter values which have these same population expansion factors in epochs 1 and 2, while the dashed red curves are the combinations of parameter values which are higher and lower by 25% compared to the true population expansion factors in each epoch. The inferred parameter combinations reflect the population expansion factors very well.

We also applied the coalescent simulation-based method of Excoffier et al. (2013) to the simulated datasets used in Figures 2–5, the results of which are shown in Figure 6. Since we are working with large sample sizes here, we restricted the method of Excoffier et al. to use 200 and 500 coalescent tree simulations per likelihood function evaluation for Scenario 1 and Scenario 2, respectively, and at most 40 rounds of conditional expectation maximization (ECM cycles). Note that in their work, Excoffier et al. (2013) use 10^5^ coalescent tree simulations per likelihood function evaluation and 20–40 ECM cycles. There is substantially more bias in the estimated growth onset times and more uncertainty in the growth rates compared to our method (see Figure 2), which should decrease if one uses more coalescent tree simulations to evaluate the likelihood at each point. However, if we had used 10^5^ trees per likelihood computation as was done in Excoffier et al. (2013), the inference would have taken an estimated 21 CPU *days* per dataset for Scenario 1 and 47 CPU *days* per dataset for Scenario 2 on average. In contrast, our method takes on average 15 CPU *seconds* per dataset for Scenario 1 and 45 CPU *minutes* per dataset for Scenario 2.

**Figure 6.**
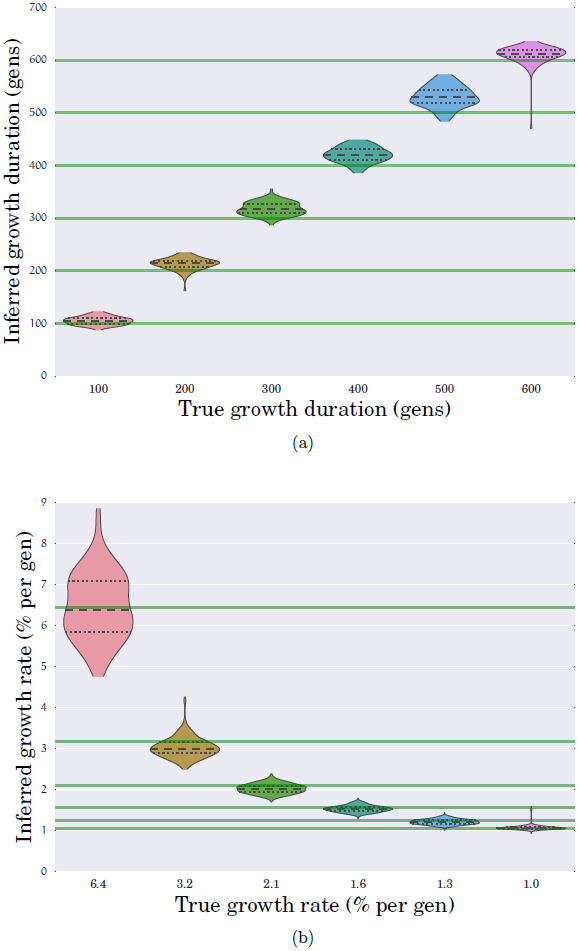
Violin plots of the (a) duration and (b) rate of exponential growth in the population size using the method of Excoffier et al. (2013) for 100 simulated datasets for each of the simulation parameter settings in Scenario 1. These are the same simulated datasets used to generate Figure 2 and Figure 3. When applying the method of Excoffier et al., due to computational reasons, we used 200 and 500 coalescent tree simulations for Scenario 1 and Scenario 2 per likelihood function estimation and limited the number of rounds of conditional expectation maximization (ECM cycles) to 40. On one of these 100 simulated datasets, their method appeared to have a runaway behavior and produced unreasonable estimates after 40 ECM cycles; this dataset was excluded from these plots.

### Per-locus estimated mutation rates

Using (8), we can compute the MLE for the mutation rate at each locus while estimating the optimal population size function parameters. The inferred mutation rates for each set of parameters in Scenario 1 and for Scenario 2 are shown in Figure 7. Since the mutation rates are estimated using the inferred demography, uncertainty in the demographic estimates will lead to uncertainty in the mutation rate estimates, as can be seen for the estimates for Scenario 2 in Figure 7. For Scenario 1 with *t*_1_ = 100 gens and *r*_1_ = 6.4% per gen, we also simulated datasets where the mutation rate at each locus is randomly chosen from the range 1.1 *×* 10^−8^ to 3.8 *×* 10^−8^ per base per gen per haploid, and then held fixed across all the simulated datasets. This is the range of mutation rates estimated from family trio data by Conrad et al. (2011). Figure 8 shows the performance of our method on simulated datasets with 100 loci each of length 10 kb, and demonstrates that our procedure can accurately recover the mutation rates.

**Figure 7.**
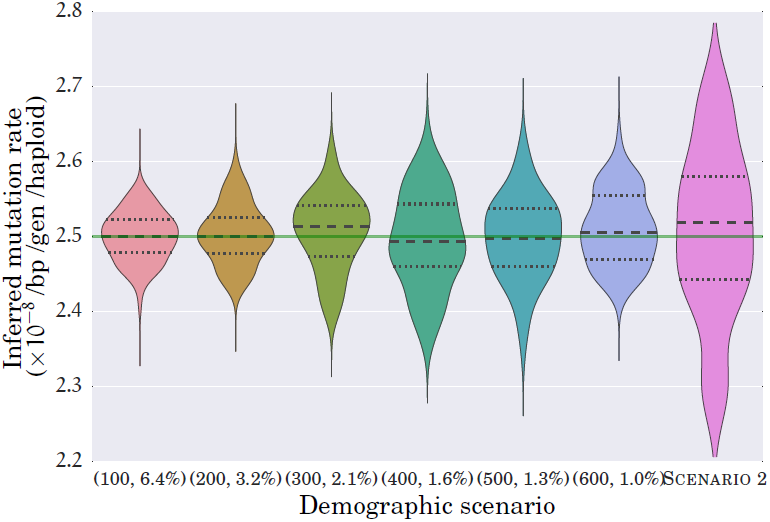
Violin plots of the inferred mutation rates for each of the six simulation parameter combinations of Scenario 1, and for Scenario 2. Each plot represents 100 simulated datasets with 100 unlinked loci of 10 kb each over 10,000 diploid individuals. All loci were simulated using a mutation rate of 2.5*×*10^−8^ per bp per gen per haploid. The uncertainty in the inferred mutation rate is significantly higher for Scenario 2 due to the higher uncertainty in the demographic parameters being simultaneously estimated (see Figure 4).

**Figure 8.**
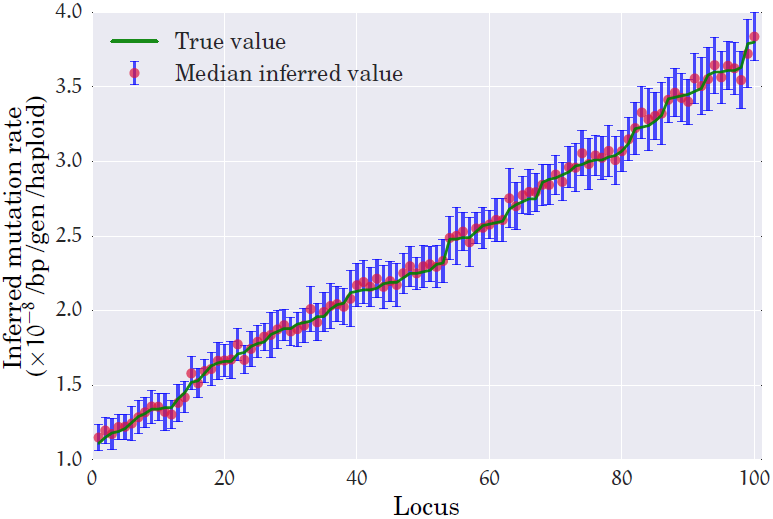
Inferred mutation rates for simulated datasets with 100 loci from 10,000 diploids under Scenario 1 with *t*_1_ = 100 and *r*_1_ = 6.4%. The mutation rates at the 100 loci were drawn randomly from the range [1.1 *×* 10^−8^, 3.8 *×* 10^−8^]. The loci are sorted in ascending order of the simulated mutation rates. The green line indicates the mutation rates used in the simulation, while the red circle and blue error bars denote the median and one standard deviation of the inferred mutation rate over 100 simulated datasets.

### Confidence intervals

Our inference algorithm described in **Method** assumes that the sites within each locus are unlinked, in which case the function *ℒ*(Φ) given in (9) is a true log-likelihood function. However, since actual genomic data (and our simulated datasets) involve non-trivial linkage within a locus, the function *ℒ*(Φ) in (9) is a composite log-likelihood function. Hence, the asymptotic confidence interval expressions in Confidence intervals will not necessarily be well calibrated. To understand this issue a bit better, we simulated datasets under Scenario 1 with *t* = 100 gens and *r*_1_ = 6.4% per gen. We generated datasets with sequence length of 10^6^ bp by simulating 10^6^*/m* unlinked loci of length *m* bp each and with recombination rate of 10^−8^ per base per gen per haploid. We did this for *m* ∈ {100, 10^3^, 10^4^} bp. By linearity of expectation, the expected total number of segregating sites is independent of the locus length *m*. Hence, for small locus lengths *m*, we expect fewer segregating sites per locus, and thus more independence between the segregating sites across the sequence. In such cases, we expect the function in (9) to be close to the true log-likelihood function. Figures 9(a) and 9(b) show asymptotic confidence intervals for the inferred growth onset times and growth rates over 100 simulated datasets. According to those figures, the asymptotic confidence interval procedure in Confidence intervals are close to an idealized confidence interval estimation procedure when the locus length *m* is shorter than 1 kb. For longer locus lengths of *m* = 10 kb, we performed a resampling block bootstrap procedure with 200 bootstrap resamples per dataset to estimate confidence intervals for the exponential population growth parameters. As shown in Figures 9(c) and 9(d), the bootstrap confidence intervals are much more faithful to an idealized confidence interval estimation procedure and are better calibrated than those produced by the asymptotic confidence interval estimation procedure.

**Figure 9.**
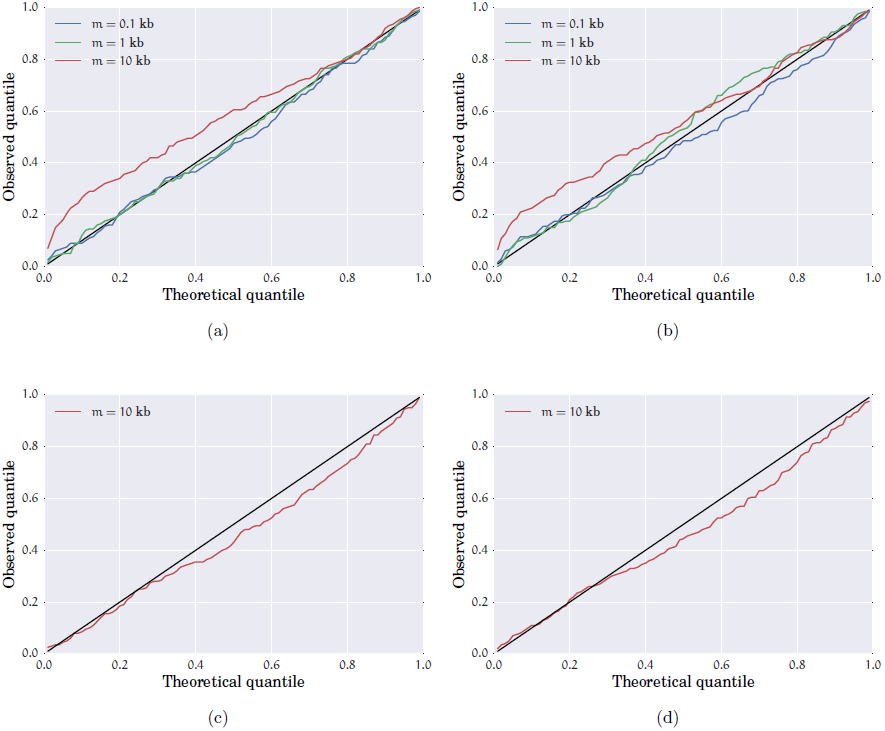
Calibration plots for (a)–(b) asymptotic, and (c)–(d) bootstrap confidence intervals of the duration and rate of exponential growth for Scenario 1 with *t*_1_ = 100 gens and *r*_1_ = 6.4% per gen for 200 simulated datasets of 10,000 diploids, each with 100 unlinked loci of length *m*. For each confidence level *α* on the *x*-axis, the *y*-axis counts the fraction of datasets where the true parameter estimates lie outside the 100(1 *− α*)% predicted confidence interval. The black lines denote the plot that would be obtained from an idealized confidence interval estimation procedure. (a) & (b): Asymptotic confidence interval calibration plots for the inferred (a) duration and (b) rate of exponential growth. As the locus length *m* increases, linkage disequilibrium causes the composite log-likelihood approximation in (9) to become increasingly inaccurate, thus leading to poorly calibrated asymptotic confidence intervals for *m* = 10 kb. (c) & (d): Bootstrap confidence interval calibration plots using 200 bootstrap replicates per simulated dataset for the inferred (c) duration and (d) rate of exponential growth. The bootstrap confidence intervals are much better calibrated than those produced by the asymptotic confidence interval estimation procedure.

### Application to real data I: Neutral regions

The dataset of Gazave et al. (2014) consists of 15 carefully curated loci from 500 individuals of European ancestry that were sequenced at a high coverage depth of 295X. These loci were chosen to be distant from known or potential coding regions, as well as regions believed to be under selection. These 15 loci contain 1,746 segregating sites that were sequenced in at least 450 individuals. Gazave et al. employed coalescent simulations to fit several demographic models incorporating recent exponential population growth to this dataset. In their models, they assumed that the ancient European demography has two population bottlenecks that were inferred by Keinan et al. (2007). Incidentally, these are also the same bottlenecks that were used in our simulation study (Scenarios 1 and 2) described above. Gazave et al.’s best-fit model (Model II) had a growth rate of 3.38% per generation starting about 140 generations in the past.

We applied our inference program to fit a model of exponential growth fixing the ancient bottlenecks to that inferred by Keinan et al. (2007). We inferred the following three parameters: the rate and the onset time of recent exponential growth, and the population size just before the onset of exponential growth. We inferred that the population grew exponentially at a rate of 3.89% per generation starting 130 generations in the past, resulting in a present effective population size of about 820 thousand individuals. Figure 10 shows the inferred demographic model, while Table 1 summarizes the point estimates and 95% confidence interval for the inferred parameter values. The confidence intervals for the demographic parameters were generated using 1000 block bootstrap resamples. These confidence intervals have significant overlap with those estimated by Gazave et al. in their best-fit model.

**Figure 10.**
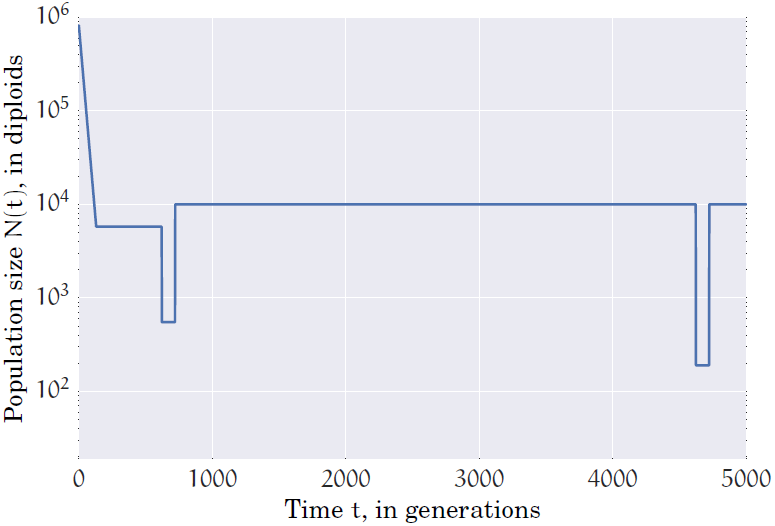
Demographic model inferred by our program on the neutral regions dataset of Gazave et al. (2014). We inferred a model with three parameters: the rate and onset of recent exponential growth, and the population size just before the onset of exponential growth. The ancient bottlenecks were fixed to those inferred by Keinan et al. (2007).

**Table 1.** Point estimates and 95% confidence intervals for the demographic parameters inferred from the neutral regions dataset of Gazave et al. (2014).

| Parameter | Point estimate | 95% confidence interval |
| --- | --- | --- |
| Population size before growth onset (diploids) | 5,769 | (4,802, 8,785) |
| Growth rate (% per gen) | 3.89% | (2.97%, 6.96%) |
| Growth onset time (gens) | 129.8 | (99.2, 162.6) |
| Population size after growth (diploids) | 818,786 | (492,068, 6,527,975) |

### Application to real data II: Exome-sequencing

Nelson et al. (2012) sequenced over 14,000 individuals from several case-control studies at 202 coding regions that are of interest for drug targeting. This dataset includes the SFS of a sample of 11,000 individuals of European ancestry containing *≈* 2,600 segregating sites among *≈* 43,000 four-fold degenerate sites in 185 genes (Nelson et al. 2012, Database S3). Using the demographic estimates of Schaffner et al. (2005) for modeling the ancient demography of the CEU population, Nelson et al. employed the coalescent simulation-based approach of Coventry et al. (2010) to fit an epoch of recent exponential growth to their data. They estimated that the effective population size of the CEU subpopulation expanded from 7,700 individuals 375 generations ago to about 4 millions individuals at the present time at a rate of about 1.68% per generation.

We also applied our method to this dataset to infer an epoch of recent exponential growth while fixing the ancient demographic model to that estimated by Schaffner et al. (2005). We inferred an epoch of exponential growth lasting 372 generations with a growth rate of 1.50% per generation. This results in a present effective population size of approximately 1.9 million individuals. We computed empirical confidence intervals for these parameter estimates using a resampling block bootstrap procedure with 1000 bootstrap resamples. Our point estimates and 95% confidence intervals for the demographic parameters are summarized in Table 2 with the inferred effective population size function shown in Figure 11. There are several reasons for the difference in demographic estimates between our method and that of Nelson et al.’s. First, the coalescent simulations of Nelson et al. were performed assuming that all sites within a locus are completely linked, while we make the opposite extreme assumption that all sites are freely recombining and evolve independently. Second, Nelson et al. used 400 simulated coalescent trees to approximate the likelihood for each combination of demographic parameters, resulting in a noisy likelihood surface. Our method, in contrast, uses exact computation to determine the expected SFS for a given demographic model. Third, they computed the log-likelihood on a discretized grid of demographic parameters, which might result in their procedure being far from the maximum likelihood estimate if the log-likelihood function is fairly flat. Our approach mitigates this problem by employing sophisticated gradient-based algorithms that can adapt their step size to the likelihood landscape.

**Figure 11.**
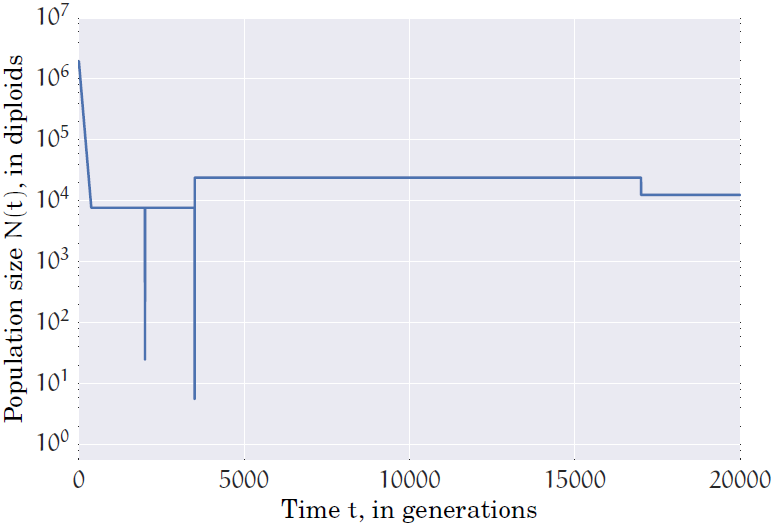
Demographic model of recent population expansion inferred by our program on the CEU exome-sequencing dataset of Nelson et al. (2012). The ancient demographic model before the epoch of exponential growth was fixed to that inferred by Schaffner et al. (2005). The two population bottlenecks 2,000 and 3,500 generations in the past correspond to the European-Asian population split and the out-of-Africa bottleneck, respectively. Our method infers an epoch of exponential population growth beginning 372 generations in the past resulting in the effective population size expanding from 7,700 individuals to 1.9 million individuals at a rate of 1.5% per generation.

**Table 2.**
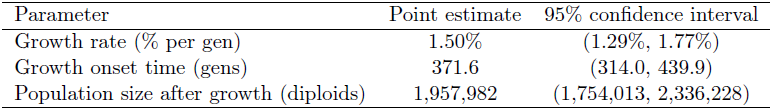
Point estimates and 95% confidence intervals for the demographic parameters inferred from the exome-sequencing dataset of Nelson et al. (2012).

We also estimated the mutation rate at each locus using the Watterson estimator given in (8). Figure 12 shows our mutation rate estimates and 95% confidence intervals, and compares them to those reported by Nelson et al. Our estimates are close to those of Nelson et al. while being systematically larger. This systematic difference can be explained by noting that our inferred population growth rate is slower than that inferred by Nelson et al., resulting in a higher mutation rate being needed to explain the same number of rare variants.

**Figure 12.**
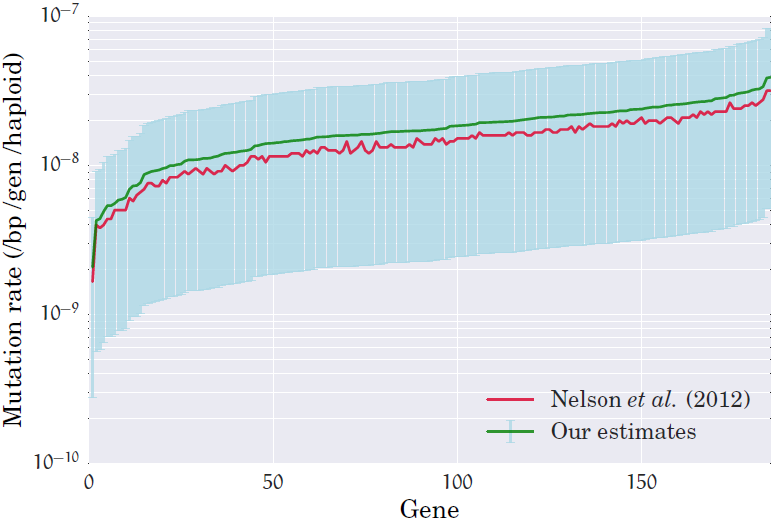
Mutation rates inferred by our method for each of the 185 genes in the exome-sequencing dataset of Nelson et al. (2012). The green line connects our point estimates for the mutation rate while the light blue lines denote 95% confidence intervals that were constructed by a resampling block bootstrap procedure with 1000 bootstrap samples. The red line connects the point estimates of the mutation rate inferred by Nelson et al. (2012). While the mutation rates estimated by our method and that of Nelson et al. are very close to each other, the mutation rates estimated by our method is systematically higher at each locus owing to the lower population expansion rate inferred by our method.

## Discussion

Aside from being of historical interest, demography is important to study because it influences genetic variation, and understanding the intricate interplay between natural selection, genetic drift, and demography is a key aim in population genomics. For example, the human census population has expanded more than 1000-fold in the last 400 generations (Keinan and Clark 2012), resulting in a state that is profoundly out of equilibrium with respect to genetic variation. Recently, there has been much interest in studying the consequences of such rapid expansion on mutation load and the genetic architecture of complex traits (Gazave et al. 2013; Simons et al. 2014; Lohmueller 2013).

Beyond developing more accurate null models of neutral evolution in order to identify genomic regions subject to natural selection (Boyko et al. 2008; Lohmueller et al. 2008; Williamson et al. 2005), the problem of inferring demography from genomic data has several other important applications. In particular, the population demography is needed to correct for spurious genotype-phenotype associations in genome-wide association studies due to hidden population substructure (Campbell et al. 2005; Clayton et al. 2005; Marchini et al. 2004); to date historical population splits, migrations, admixture and introgression events (Gravel et al. 2011; Li and Durbin 2011; Lukić and Hey 2012; Sankararaman et al. 2012); to compute random match probabilities accurately in forensic applications (Balding and Nichols 1997; Graham et al. 2000); and so on.

In this paper, we developed a method that can leverage genomic variation data from samples involving tens of thousands of individuals to infer details of very recent changes in the effective population size. Our method is based on efficient exact computation of the expected SFS of a sample, and scales well to the large sample sizes that are common in genomic sequencing studies today. Taking advantage of the fact that a wide class of parametric demographic models are uniquely determined by the SFS of even moderate size samples, we focused on inferring piecewise-exponential models of recent population size changes. Our inference procedure was derived under an assumption that different sites evolved in an independent fashion.

Our approach suggests several directions for future research. It would be interesting to investigate other families of parametric population size functions that might better fit human genetic data. For example, Reppell et al. (2014) studied more general population growth models which allow for super- and subexponential growth through the incorporation of an acceleration parameter. Such richer parametric families might allow one to fit demographic models with fewer number of pieces to the data. It is easy to extend our method to such parametric families, since the only dependence on the functional form of the population size function is in the integral expressions given in the Supplemental material.

It would also be useful to extend the analytic coalescent theory of the SFS to more realistic demographic models that incorporate multiple populations with migrations and time-varying population sizes. This would enable one to develop demographic inference algorithms that are potentially more efficient than coalescent simulation-based methods or numerical diffusion-based methods. A natural extension to the SFS which captures variation in samples from multiple subpopulations is the joint population SFS which counts the number of segregating sites in the sample as a function of the sample allele frequency in each subpopulation. Chen (2012) has developed analytic expressions for the expected joint population SFS of multiple subpopulations when the migrations are instantaneous exchanges of individuals between subpopulations. Even more fundamentally, it would be interesting to characterize the statistical identifiability of complex demographic models from the joint population SFS of random samples.

Another direction for research is understanding how the uncertainties in the inference of different parameters are related to each other. For example, the violin plots in Figure 2 show that when there is more uncertainty in the inference of the exponential growth rate, there is less uncertainty in the inference of the growth duration, and vice versa. A similar observation can be made from Figure 4 for the inference of the exponential growth parameters of the two epochs in Scenario 2. It would be interesting to understand if there is a fundamental quantifiable uncertainty relation that is independent of the inference algorithm, or if this behavior is specific to our inference procedure.

## Acknowledgements

A.B. thanks Andrew Chan for helpful discussions at the initial stages of this work. We are grateful to John Novembre and Darren Kessner for sharing their demographic estimates on the exome-sequencing dataset, and to Alon Keinan and Li Ma for sharing the SFS of the neutral regions dataset. This research is supported in part by an NIH grant R01-GM094402, a Packard Fellowship for Science and Engineering, and a Simons-Berkeley Research Fellowship.

## References

Balding, D. J. and Nichols, R. A., 1997. Significant genetic correlations among Caucasians at forensic DNA loci. Heredity, 78(6).

Bhaskar, A. and Song, Y. S., 2013. The identifiability of piecewise demographic models from the sample frequency spectrum. arXiv preprint arXiv:1309.5056.

Boyko, A. R., Williamson, S. H., Indap, A. R., Degenhardt, J. D., Hernandez, R. D., Lohmueller, K. E., Adams, M. D., Schmidt, S., Sninsky, J. J., Sunyaev, S. R., et al., 2008. Assessing the evolutionary impact of amino acid mutations in the human genome. PLoS Genetics, 4(5):e1000083.

Campbell, C. D., Ogburn, E. L., Lunetta, K. L., Lyon, H. N., Freedman, M. L., Groop, L. C., Altshuler, D., Ardlie, K. G., and Hirschhorn, J. N., 2005. Demonstrating stratification in a European American population. Nature Genetics, 37(8):868–872.

Chen, H., 2012. The joint allele frequency spectrum of multiple populations: A coalescent theory approach. Theoretical Population Biology, 81(2):179–195.

Clayton, D. G., Walker, N. M., Smyth, D. J., Pask, R., Cooper, J. D., Maier, L. M., Smink, L. J., Lam, A. C., Ovington, N. R., Stevens, H. E., et al., 2005. Population structure, differential bias and genomic control in a large-scale, case-control association study. Nature Genetics, 37(11):1243–1246.

Conrad, D., Keebler, J., DePristo, M., Lindsay, S., Zhang, Y., Casals, F., Idaghdour, Y., Hartl, C., Torroja, C., Garimella, K., et al., 2011. Variation in genome-wide mutation rates within and between human families. Nature, 201:1.

Coventry, A., Bull-Otterson, L. M., Liu, X., Clark, A. G., Maxwell, T. J., Crosby, J., Hixson, J. E., Rea, T. J., Muzny, D. M., Lewis, L. R., et al., 2010. Deep resequencing reveals excess rare recent variants consistent with explosive population growth. Nature Communications, 1:131.

Efron, B. and Tibshirani, R., 1986. Bootstrap methods for standard errors, confidence intervals, and other measures of statistical accuracy. Statistical science, 1(1):54–75.

Ewens, W., 2004. Mathematical Population Genetics: I. Theoretical Introduction. Springer, 2nd edition.

Excoffier, L., Dupanloup, I., Huerta-Sánchez, E., Sousa, V. C., and Foll, M., 2013. Robust demographic inference from genomic and SNP data. PLoS Genetics, 9(10):e1003905.

Fu, W., O’Connor, T. D., Jun, G., Kang, H. M., Abecasis, G., Leal, S. M., Gabriel, S., Altshuler, D., Shendure, J., Nickerson, D. A., et al., 2012. Analysis of 6,515 exomes reveals the recent origin of most human protein-coding variants. Nature, 493:216–220.

Fu, Y.-X., 1995. Statistical properties of segregating sites. Theoretical Population Biology, 48:172–197.

Gazave, E., Chang, D., Clark, A. G., and Keinan, A., 2013. Population growth inflates the per-individual number of deleterious mutations and reduces their mean effect. Genetics, 195(3):969–978.

Gazave, E., Ma, L., Chang, D., Coventry, A., Gao, F., Muzny, D., Boerwinkle, E., Gibbs, R. A., Sing, C. F., Clark, A. G., et al., 2014. Neutral genomic regions refine models of recent rapid human population growth. Proceedings of the National Academy of Sciences, 111(2):757–762.

Graham, J., Curran, J., and Weir, B., 2000. Conditional genotypic probabilities for microsatellite loci. Genetics, 155(4):1973–1980.

Gravel, S., Henn, B. M., Gutenkunst, R. N., Indap, A. R., Marth, G. T., Clark, A. G., Yu, F., Gibbs, R. A., Bustamante, C. D., Altshuler, D. L., et al., 2011. Demographic history and rare allele sharing among human populations. Proceedings of the National Academy of Sciences, 108(29):11983–11988.

Griewank, A. and Corliss, G. F., 1991. Automatic differentiation of algorithms: theory, implementation, and application. Society for industrial and Applied Mathematics Philadelphia, PA.

Gutenkunst, R. N., Hernandez, R. D., Williamson, S. H., and Bustamante, C. D., 2009. Inferring the joint demographic history of multiple populations from multidimensional SNP frequency data. PLoS Genetics, 5(10):e1000695.

Harris, K. and Nielsen, R., 2013. Inferring demographic history from a spectrum of shared haplotype lengths. PLoS Genetics, 9(6):e1003521.

Hudson, R., 2002. Generating samples under a Wright–Fisher neutral model of genetic variation. Bioinformatics, 18(2):337–338.

Keinan, A. and Clark, A. G., 2012. Recent explosive human population growth has resulted in an excess of rare genetic variants. Science, 336(6082):740–743.

Keinan, A., Mullikin, J. C., Patterson, N., and Reich, D., 2007. Measurement of the human allele frequency spectrum demonstrates greater genetic drift in East Asians than in Europeans. Nature Genetics, 39(10):1251–1255.

Li, H. and Durbin, R., 2011. Inference of human population history from individual whole-genome sequences. Nature, 475(7357):493–496.

Lohmueller, K. E., 2013. The impact of population demography and selection on the genetic architecture of complex traits. arXiv preprint arXiv:1306.5261.

Lohmueller, K. E., Indap, A. R., Schmidt, S., Boyko, A. R., Hernandez, R. D., Hubisz, M. J., Sninsky, J. J., White, T. J., Sunyaev, S. R., Nielsen, R., et al., 2008. Proportionally more deleterious genetic variation in European than in African populations. Nature, 451(7181):994–997.

Lukić, S. and Hey, J., 2012. Demographic inference using spectral methods on SNP data, with an analysis of the human out-of-Africa expansion. Genetics, 192(2):619–639.

Lukić, S., Hey, J., and Chen, K., 2011. Non-equilibrium allele frequency spectra via spectral methods. Theoretical Population Biology, 79(4):203–219.

Marchini, J., Cardon, L. R., Phillips, M. S., and Donnelly, P., 2004. The effects of human population structure on large genetic association studies. Nature Genetics, 36(5):512–517.

Marth, G., Czabarka, E., Murvai, J., and Sherry, S., 2004. The allele frequency spectrum in genome-wide human variation data reveals signals of differential demographic history in three large world populations. Genetics, 166(1):351–372.

Myers, S., Fefferman, C., and Patterson, N., 2008. Can one learn history from the allelic spectrum? Theoretical Population Biology, 73(3):342–348.

Nelson, M. R., Wegmann, D., Ehm, M. G., Kessner, D., Jean, P. S., Verzilli, C., Shen, J., Tang, Z., Bacanu, S.-A., Fraser, D., et al., 2012. An abundance of rare functional variants in 202 drug target genes sequenced in 14,002 people. Science, 337(6090):100–104.

Nielsen, R., 2000. Estimation of population parameters and recombination rates from single nucleotide polymorphisms. Genetics, 154(2):931–942.

Palamara, P. F., Lencz, T., Darvasi, A., and Pe’er, I., 2012. Length distributions of identity by descent reveal fine-scale demographic history. American Journal of Human Genetics, 91(5):809–822.

Polanski, A., Bobrowski, A., and Kimmel, M., 2003. A note on distributions of times to coalescence, under time-dependent population size. Theoretical Population Biology, 63(1):33–40.

Polanski, A. and Kimmel, M., 2003. New explicit expressions for relative frequencies of single-nucleotide polymorphisms with application to statistical inference on population growth. Genetics, 165(1):427–436.

Reppell, M., Boehnke, M., and Zöllner, S., 2014. The impact of accelerating, faster than exponential population growth on genetic variation. Genetics, 196(3):819–828.

Sankararaman, S., Patterson, N., Li, H., Pääbo, S., and Reich, D., 2012. The date of interbreeding between Neandertals and modern humans. PLoS Genetics, 8(10):e1002947.

Sawyer, S. A. and Hartl, D. L., 1992. Population genetics of polymorphism and divergence. Genetics, 132(4):1161–76.

Schaffner, S., Foo, C., Gabriel, S., Reich, D., Daly, M., and Altshuler, D., 2005. Calibrating a coalescent simulation of human genome sequence variation. Genome Research, 15(11):1576–1583.

Sheehan, S., Harris, K., and Song, Y. S., 2013. Estimating variable effective population sizes from multiple genomes: A sequentially Markov conditional sampling distribution approach. Genetics, 194:647–662.

Simons, Y. B., Turchin, M. C., Pritchard, J. K., and Sella, G., 2014. The deleterious mutation load is insensitive to recent population history. Nat. Genet., 46:220–224.

Tennessen, J. A., Bigham, A. W., O’Connor, T. D., Fu, W., Kenny, E. E., Gravel, S., McGee, S., Do, R., Liu, X., Jun, G., et al., 2012. Evolution and functional impact of rare coding variation from deep sequencing of human exomes. Science, 337(6090):64–69.

Walther, A. and Griewank, A., 2012. Getting started with ADOL-C. In und O. Schenk, U. N., editor, Combinatorial Scientific Computing, pages 181–202. Chapman-Hall CRC Computational Science.

Watterson, G., 1975. On the number of segregating sites in genetical models without recombination. Theoretical Population Biology, 7(2):256–276.

Williamson, S. H., Hernandez, R., Fledel-Alon, A., Zhu, L., Nielsen, R., and Bustamante, C. D., 2005. Simultaneous inference of selection and population growth from patterns of variation in the human genome. Proceedings of the National Academy of Sciences, 102(22):7882–7887.

